# Bayesian multi-model-based ^13^C^15^N-metabolic flux analysis quantifies carbon-nitrogen metabolism in mycobacteria

**DOI:** 10.1101/2022.03.08.483448

**Authors:** Khushboo Borah, Martin Beyß, Ye Xu, Jim Barber, Catia Costa, Jane Newcombe, Axel Theorell, Melanie J Bailey, Dany JV Beste, Johnjoe McFadden, Katharina Nöh

## Abstract

Metabolic flux is the final output of cellular regulation and has been extensively studied for carbon but much less is known about nitrogen, which is another important building block for living organisms. For the pathogen *Mycobacterium tuberculosis* (Mtb), this is particularly important in informing the development of effective drugs targeting Mtb’s metabolism. Here we performed ^13^C^15^N dual isotopic labelling of mycobacterial steady state cultures and quantified intracellular carbon-nitrogen (CN) and nitrogen (N) fluxes in addition to carbon (C) fluxes and inferred their reaction bidirectionalities. The combination of ^13^C^15^N-MFA with a Bayesian multi-model approach allowed us to resolve C and N fluxes simultaneously which was not possible with classical ^13^C-MFA. We quantified CN fluxes for amino acid and, for the first time, nucleotide biosynthesis. Our analysis identified glutamate as the central CN and N node in mycobacteria, and improved resolution of the anaplerotic node. Our study describes a powerful platform to measure carbon and nitrogen metabolism in any biological system with statistical rigor.

## Introduction

In recent decades, a great deal of progress has been made in unravelling the complexity of intracellular metabolism in microbial, animal and plant cells by measuring metabolic fluxes through the reactions that constitute central metabolism. The state-of-the-art technique is ^13^C-Metabolic Flux Analysis (MFA) in which cells, at metabolic steady-state, are fed a mixture of ^12^C and ^13^C-labelled substrates that are incorporated into the central carbon (C) metabolism to yield stable end products, such as the proteinogenic amino acids. The method infers *in vivo* metabolic reaction rates (fluxes) by using a system-wide biochemical reaction model that tracks C atom rearrangements throughout the metabolic pathways, and by fitting these fluxes to the emerging labelling patterns (typically isotopically ^12^C and ^13^C labelled fractional enrichments measured by mass spectrometry (MS) or nuclear magnetic resonance (NMR)) (*1*–*4*). ^13^C-MFA resolves the activity of biochemical reactions through computational modelling which can differentiate between parallel pathways and determine bidirectional fluxes (mass exchange of reactions that proceed forwards and backwards at the same time) (*5*, *6*).

Besides central C metabolism, nitrogen (N) metabolism plays a key role, not only in amino acid and nucleotide metabolism, but also in the synthesis of many cofactors (*7*). In many microbes including the pathogenic *Mycobacterium tuberculosis*, nitrate acts as a terminal oxygen acceptor in addition to molecular oxygen during hypoxic respiration (*8*). Although N fixation and assimilation play a key role in medical research, agriculture and biotechnology, quantitative insights into N metabolism are currently limited. Consequently, only a few drugs have been developed that target N metabolism (*9*). The progress in quantifying N metabolism has been challenging, mostly because there is limited information derived from the isotopic labelling profiles of N versus C atoms. The equivalent of ^13^C-MFA, namely ^15^N-MFA therefore needs to involve time-resolved labeling data and the measurement of intracellular intermediate metabolite concentrations (pool sizes) to deploy the isotopically non-stationary (INST) MFA framework (10–14). This technique was utilized to study ammonium assimilation in *Corynebacterium glutamicum* by quantifying central N fluxes *in vivo* (*12*).

C and N metabolism are interdependent in all organisms (*13*, *14*). For instance, the tricarboxylic acid cycle (TCA), glycolysis and pentose phosphate pathway (PPP), which are C based, synthesize amino acids and nucleotides. Biosynthesis of amino acids primarily involves addition of N to the C backbone, requiring reductants and energy, generated primarily from C metabolism (Fig. 1). To understand C and N co-assimilation, a systems-based analysis is required which includes both central C and N metabolism. The information about ^13^C labelling enrichments of the intermediates in amino acid biosynthesis alone is limited, because such measurements cannot resolve alternative pathways in which no C scrambling occurs, as in the case of arginine biosynthesis pathway (Fig.1, inset). The key for quantification of C and N fluxes simultaneously is to administer ^13^C and ^15^N isotopic tracers. Using ^13^C_6_^15^N_4_ labelled arginine as the tracer, this co-labelling strategy was implemented to study arginine metabolism in *Kluyveromyces lactis* (*15*); this study showed that ^13^C incorporation into the amino acids proceeded at a slower timescale than the ^15^N-label. The time-deconvolution of C and N label incorporation allowed approximation of C and N fluxes using a “staggered” INST ^13^C-MFA and ^15^N-MFA approach. However, the efficacy of this approach critically depends on the validity of the underlying timescale separation, which is induced by the efficacious choice of C- and N-sources. Here we have established an approach that simultaneously quantifies C and N metabolic fluxes independently of the co-labelling strategy applied. Our ^13^C^15^N-MFA platform specifically tracks C and N atom interconversions throughout the entire metabolic network, without the need to acquire the pool sizes of metabolic intermediates or the deconvolution of ^13^C and ^15^N isotopologues using time-deconvolution or specialized analytical measurement platforms. We demonstrate the versatility of this platform by quantifying intracellular C and N fluxes of the vaccine strain of mycobacteria *Mycobacterium bovis* BCG, as a model for one of the world’s most important pathogens,*Mycobacterium tuberculosis* (Mtb) that causes tuberculosis (TB).

**Fig. 1.**
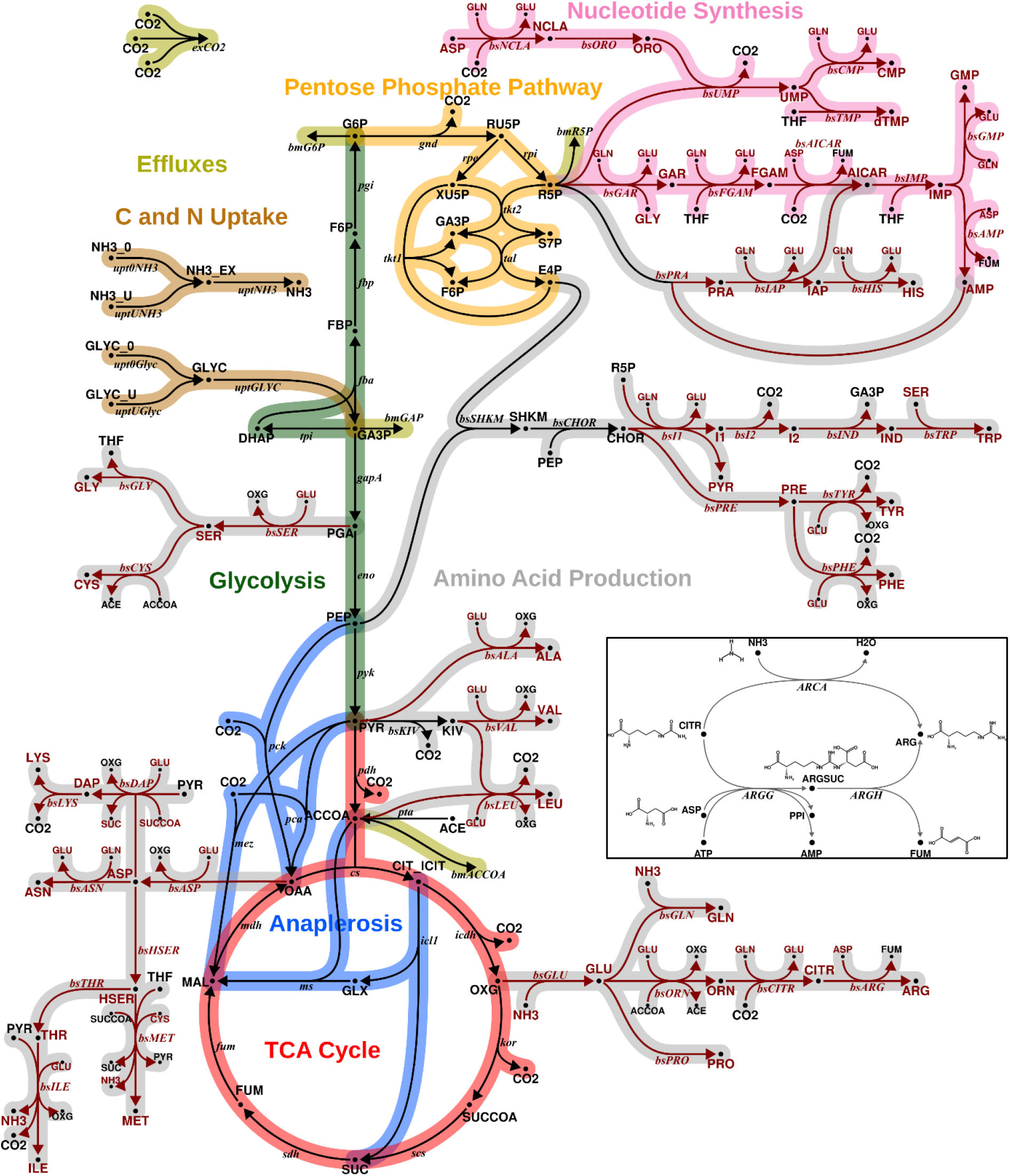
Metabolic network showing carbon and nitrogen metabolism. Pathways of carbon and nitrogen metabolism include glycolysis (EMP), pentose phosphate pathway (PPP), tricarboxylic acid (TCA) cycle, and anaplerotic reactions (ANA). Nitrogen source ammonium; carbon source glycerol. Reactions and metabolites involving nitrogen are shown in red. The inset shows the last bifurcated step of the arginine biosynthesis, according to the genome-scale metabolic model sMTB2.0 (*63*). Citrulline is aminated either by free nitrogen to form arginine (arginine deiminase, *ARCA*), or aspartate is acting as nitrogen donor and arginine is formed via a two-step reaction with the intermediate argininosuccinate (argininosuccinate synthase (*ARGG*) and argininosuccinate lyase (*ARGH*)). Because the carbon backbone is the same for both branches, ^13^C labelling alone is not able to resolve the fluxes of either of these pathways.

TB is one of the leading causes of human mortality from a single infectious agent that kills over a million people every year (*16*). Drug resistance is a major problem affecting TB therapy (*17*, *18*), so new drugs are urgently needed. Measurement of N metabolic fluxes has the potential to identify novel anti-TB drug targets, but the current progress is hampered by the limitation of tools and technology to measure N along with C fluxes *in vivo*. We previously developed ^13^C-flux spectral analysis (FSA) and ^15^N-flux spectral ratio analysis (FSRA) for identifying the probable spectrum of C and N substrates in Mtb *ex vivo* (*19*, *20*). Using FSRA, we found aspartate, glutamate, and glutamine to be the primary nitrogen sources for intracellular Mtb (*20*). Although ^13^C-FSA and ^15^N-FSRA provided qualitative conclusions about C and N sources, the available measurements did not allow for the flux quantifications. Multiple studies have successfully measured C fluxes in Mtb growing as batch cultures (*21*–*23*). We also applied ^13^C-MFA to quantify intracellular C fluxes of Mtb and BCG during slow and fast growth in a chemostat (*22*). Here using C and N isotopic co-labeling, metabolic modeling, and Bayesian statistics we resolved the central C and N co-metabolism with an increased resolution.

The measurement of N metabolic fluxes, simultaneously with the C fluxes in a principled, system-wide manner has not been attempted in Mtb or in any organism. The simultaneous measurement of C and N fluxes requires constructing an enlarged metabolic reaction model that describes both central C and N metabolism along with multi-atom transitions. As a result, the number of unknown flux parameters to be inferred from the experimental measurements, increases significantly. The increase in dimensionality stems primarily from the reaction steps for CN co-assimilation (mainly reactions catalyzed by transaminases) as these reactions must be modelled bidirectional to describe the co-labeling enrichments. In this situation, employing the standard best-fit approach commonly used in ^13^C-MFA is prone to overfitting (*24*, *25*). This problem is exacerbated when measurements cannot distinguish between labelling contributions stemming from ^13^C and ^15^N tracers, such as measurements obtained from single quadrupole MS instruments with insufficient mass resolution. To overcome the impediments of current single-model ^13^C-MFA approaches, we used a statistically rigorous multi-model inference approach (*26*), which we here generalize to ^13^C^15^N-MFA. This multi-model-based ^13^C^15^N-MFA platform for analyzing co-labeling datasets enabled us to measure intracellular metabolic fluxes for the central C and N metabolism in *M. bovis* BCG under steady-state conditions.

## Results

### Roadmap for Bayesian multi-model ^13^C^15^N-metabolic flux analysis

The ^13^C^15^N-MFA co-labelling general workflow is summarized in Fig. 2. Cultivation experiments are performed under metabolic (pseudo) steady state conditions, in a C or N limited chemostat. Steady state cultures are switched to media containing ^13^C- and ^15^N-labelled substrates, and samples are drawn after an isotopic steady state labelling is achieved for both C and N. The samples are then analyzed by MS, providing mass isotopomer distributions (MIDs) (*27*). Although we focus on MS as mainstream analytics, the workflow is equally valid for NMR delivering heteronuclear NMR moieties (*28*), or a combination of MS and NMR measurements. In terms of mass shifts, low-resolution gas-chromatography (GC-MS) and liquid-chromatography mass spectrometry (LC-MS) are often not sufficiently sensitive to distinguish between ^13^C and ^15^N isotopomers (*29*), resulting in convoluted univariate (^13^C^15^N) MIDs (*30*). For example, in lysine with six C (#*C* = 6) and two N (#*N* = 2) atoms, #*C* + #*N* + 1 = 9 univariate MIDs exist. More advanced analytical platforms such as high-resolution LC-MS (orbitrap) (*27*), FT-ICR-MS (*31*), multi-reflection time-of-flight (ToF) MS or tailored derivatization approaches combined with two-stage LC-MS (*29*) can distinguish between ^13^C and ^15^N isotopomers, providing multivariate MIDs. In the lysine example, there are (#*C* + 1) × (#*N* + 1) = 21 multivariate mass isotopomers. Ultra-high-resolution orbitrap and FT-ICR analyzers allow resolving the full spectrum. In practice, however, those analytical platforms operate at a trade-off between resolving power (~65,000 fwhm needed to separate ^13^C and ^15^N mass isotopomers) and acquisition accuracy. Also, low intracellular metabolite concentrations often limits the precise measurements of multivariate MIDs. Therefore, we opted to measure univariate MIDs from the co-labelling experiment using a single-quadrupole GC-MS system, which is a robust analytical device for MID analysis. Our workflow is transferable and equally applicable to other analytical platforms.

**Fig. 2.**
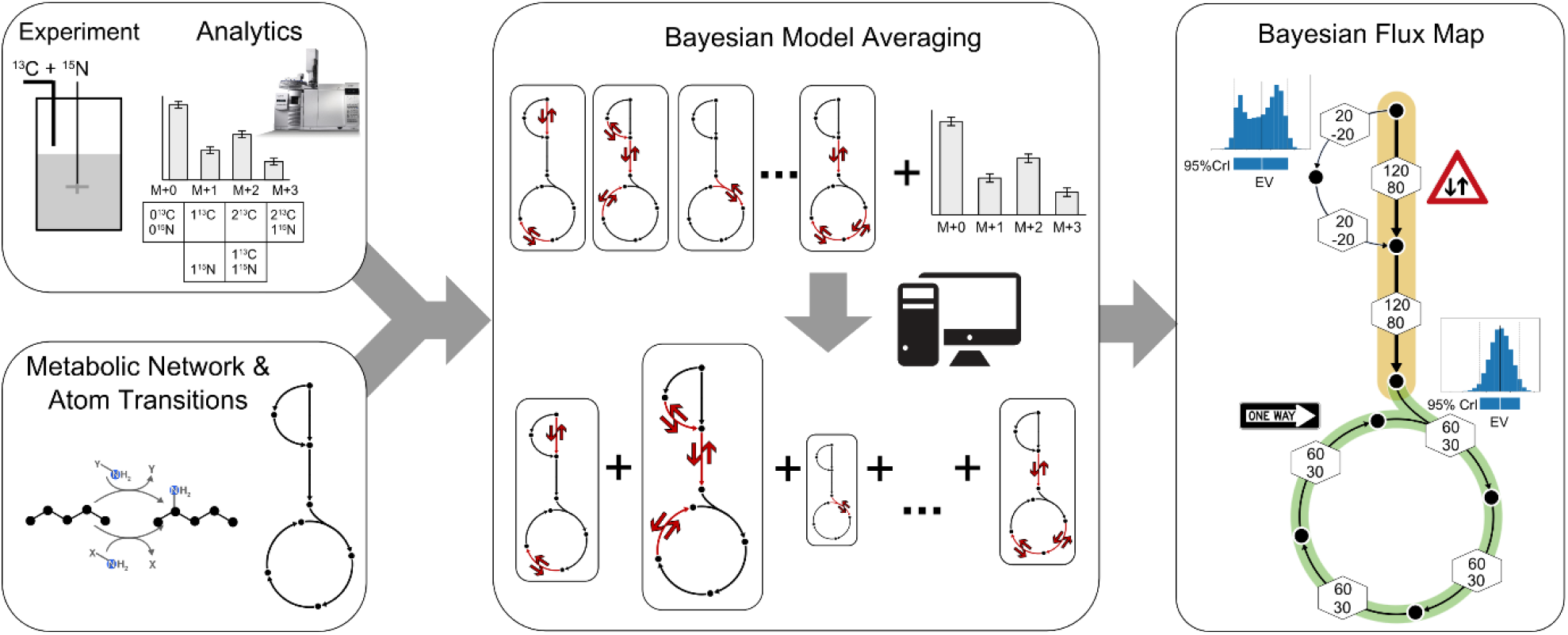
General workflow of ^13^C^15^N-MFA. Labelling data are collected from a ^13^C and ^15^N isotope co-labelling experiment, performed for a continuous culture in a chemostat setting to achieve metabolic (pseudo) steady state conditions. Cells are harvested at isotopic steady state for the analysis of intracellular metabolites using mass spectrometry along with natural abundance correction. A CN metabolic model is constructed and together with extracellular (uptake, secretion) rates and the labelling data, CN fluxes are inferred. To this end, a multi-model inference strategy using Bayesian Model Averaging (BMA) is executed. Here, any specific combination of uni- and bidirectional reactions constitutes a model, giving rise to combinatorically many possible model variants. BMA is a statistical procedure to draw inferences from the set of model variants by weighting individual model inferences based on their likelihood to explain the labelling data. The result is the Bayesian flux map that shows the resulting expected values of net fluxes, resulting from marginal posterior probability distributions, along with the probabilities of the reversible reactions to operate bidirectionally.

MIDs corrected for natural abundance, along with the extracellular rates and biomass proportions, are incorporated into a metabolic network model that precisely specifies the transition of C and N atoms throughout the intracellular reactions. Tracking N in addition to C requires not only an extension of C mappings by N mappings and the addition of the reactions of nitrogen metabolism, but also requires a refined formulation of biosynthesis reactions that are usually lumped in ^13^C-MFA (an example is shown in Fig. 1 inset). These extensions enable us to infer CN fluxes from the co-labelling data. In addition to the transition network, information on the mass exchange between intermediates of these biosynthetic reactions i.e., whether the mass flow through these reversible reactions is unidirectional or bidirectional is required (*6*). Transaminases are suspected to operate at near thermodynamic equilibrium, but quantitative evidence regarding their activity are largely missing (*32*). As such all transaminase catalyzed biochemical reactions carrying CN fluxes should be considered potentially bidirectional (*33*), implying that they are characterized by two flux parameters, a net and an exchange flux, instead of a unidirectional reaction, which is described by a net flux only. Consequently, this introduces additional challenges to identify flux parameters into the CN model, which renders the model susceptible to overfitting. A general solution for dealing with model under-determinacy in a statistically rigorous manner, while making as few assumptions as possible, has been proposed within the framework of Bayesian Model Averaging (BMA) (*34*). When applied to ^13^C^15^N-MFA, BMA determines the probability distributions of net fluxes (*v*) given the data (*D*), in the Bayesian paradigm expressed as *p*(*v*|*D*), by averaging the flux posterior probabilities over all possible models (*M_i_*, *i* = 1,… *N*), weighted by the model probability in view of the data *p*(*M_i_*|*D*):

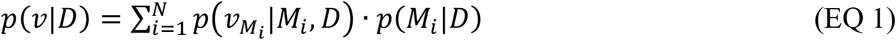

Here, models *M_i_*, are structural variants that differ in their bidirectionality setting and, hence, number of flux parameters (*v_M_i__*). Equation (EQ 1) is solved computationally by using a recently developed tailored Markov chain Monte Carlo (MCMC) approach (*26*) (see Materials and Methods for details). This results in so-called marginal posterior probability distributions for the net fluxes, as well as the probabilities of reversible reactions being uni- or bidirectional. The marginal posterior probability distributions provide credible intervals (CrI) and expected values (EV) for the net fluxes. Finally, this outcome is visually summarized in a Bayesian flux map (Fig. 2). In addition, two-dimensional marginal posterior probability distributions give insights into possibly non-linear net flux correlations. It should be noted that all traditional flux maps, including those in our previous work (*22*), instead report maximum likelihood-based point estimates and confidence intervals while flux correlations are exclusive to the Bayesian framework.

### BMA-based ^13^C^15^N-MFA validates and refines carbon fluxes determined using ^13^C-MFA in mycobacteria

We applied the ^13^C^15^N-MFA workflow to the mycobacterial model system *M. bovis* BCG. The experimental conditions were comparable to those described in Beste *et al*. 2011 (*22*). *M. bovis* BCG was cultivated in continuous culture with glycerol and ammonia as sole C and N sources respectively at a dilution rate of 0.03 h^-1^ (Table 1, Fig. S1A-C). Cultures were grown with 10% [^13^C_3_]-glycerol (GLYC) and 20% [^15^N_1_]-ammonium chloride (NH_4_Cl) until an isotopic steady state was reached, as confirmed by GC-MS of amino acids (Fig. S1C). Harvested samples were analyzed using GC-MS, providing univariate MID measurements (MIDs which do not distinguish between ^13^C and ^15^N isotopomers) of 15 proteinogenic amino acids.

**Table 1.**
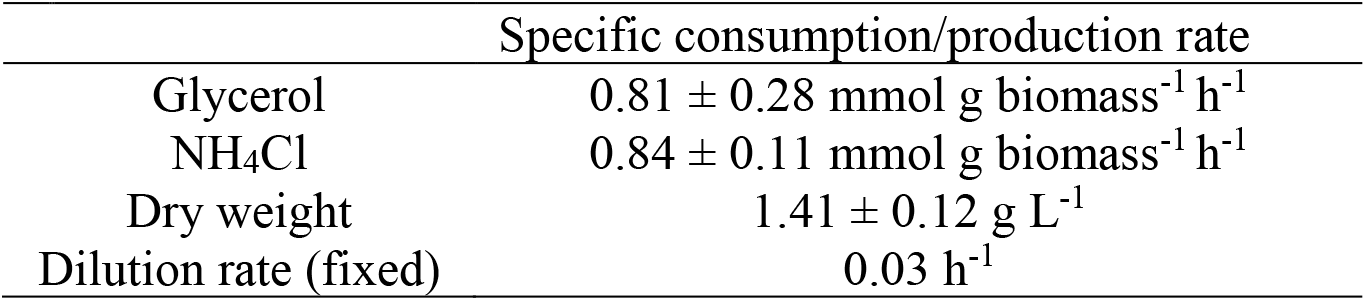
The measurements were done for chemostat cultures at metabolic and isotopic steady state. Measurements are mean ± SD from two independent chemostat cultures each with three to four technical replicates.

The resulting Bayesian flux map is shown in Fig. 3 with fluxes relative to the glycerol (GLYC) uptake rate, while absolute fluxes are given in Fig. 4 (see also supplementary Fig. S2 for the absolute net flux posterior probability distributions). As expected, the primary C metabolic route is directed from GLYC over lower glycolysis towards lipid and fatty acid synthesis (via the acetyl-CoA drain flux). While glycolytic fluxes were the highest, the TCA cycle, PPP and anaplerotic fluxes are significantly lower. The fluxes through the decarboxylating arm of the TCA cycle, oxoglutarate ferredoxin oxidoreductase (*kor*) and succinyl-CoA synthetase (*scs*) reactions are reduced; decarboxylation is bypassed using glyoxylate shunt, instantiating the GAS pathway (*22*). The net C flux distributions through the upper glycolysis, the TCA and anaplerosis for BCG growing at a growth rate (0.03h^-1^) measured using Bayesian ^13^C^15^N-MFA are very similar to those derived by traditional ^13^C-MFA in our previous study (*22*) (supplementary Fig. S3).

**Fig. 3.**
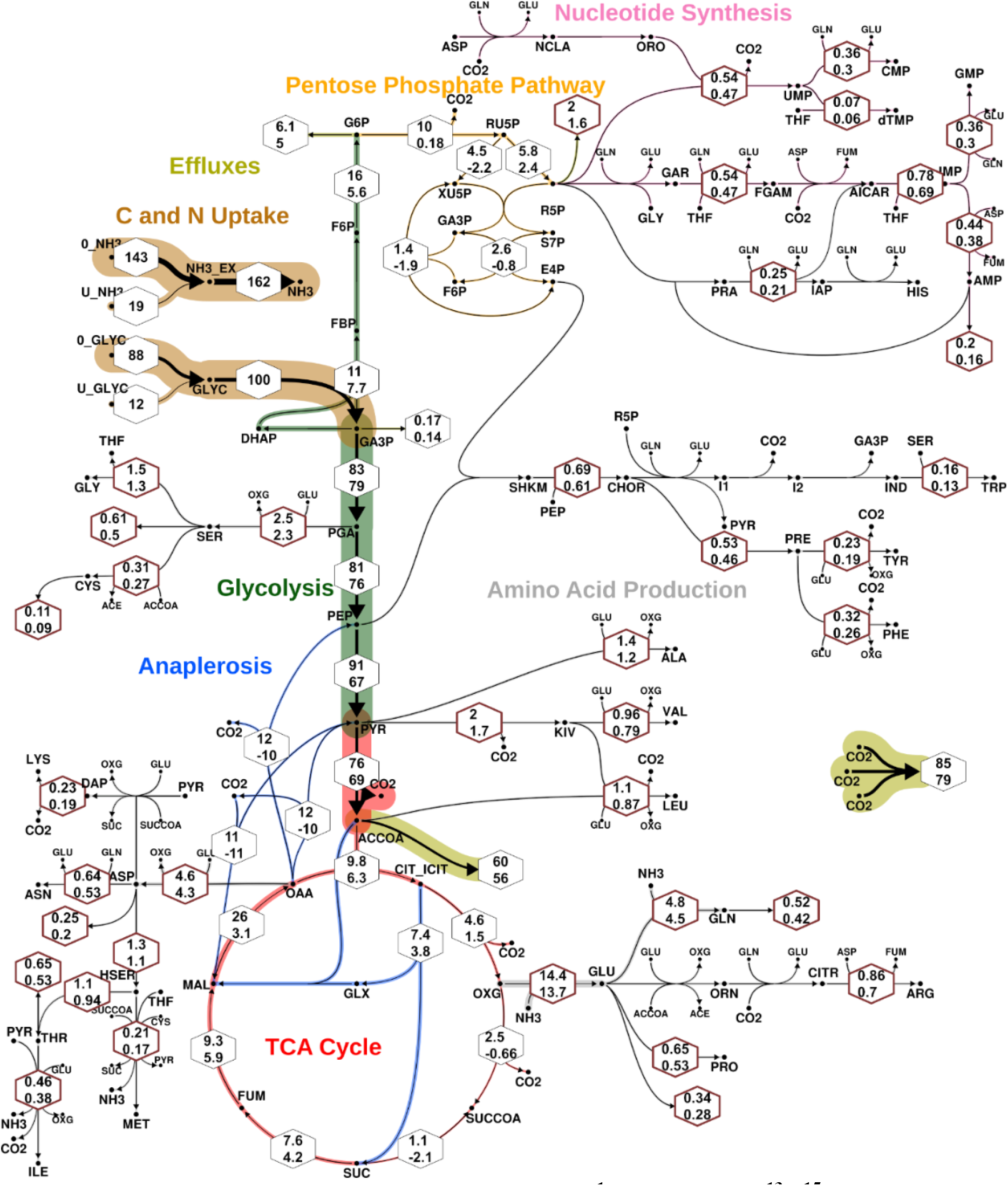
Bayesian flux map for BCG growing at 0.03 h^-1^ inferred with ^13^C^15^N-MFA. The line strengths code for the expected values (EV) of the net flux marginal posterior probability distributions (supplementary Fig. S2). Their associated 95% credible intervals (CrIs) are given in hexagons, where thin black and thick dark red borders indicate reactions that involve carbon- or nitrogen-only and mixed carbon-nitrogen transfer, respectively. Values are given relative to the glycerol uptake flux (set to 100). The associated absolute net flux CrIs are provided in Fig. 4. The nominal reaction direction, indicated by the arrowhead, is given according to a positive EV. Colour indicates pathway association of the reactions. Associated probabilities of reversible reactions being bidirectional are found in supplementary Fig. S6.

**Fig. 4.**
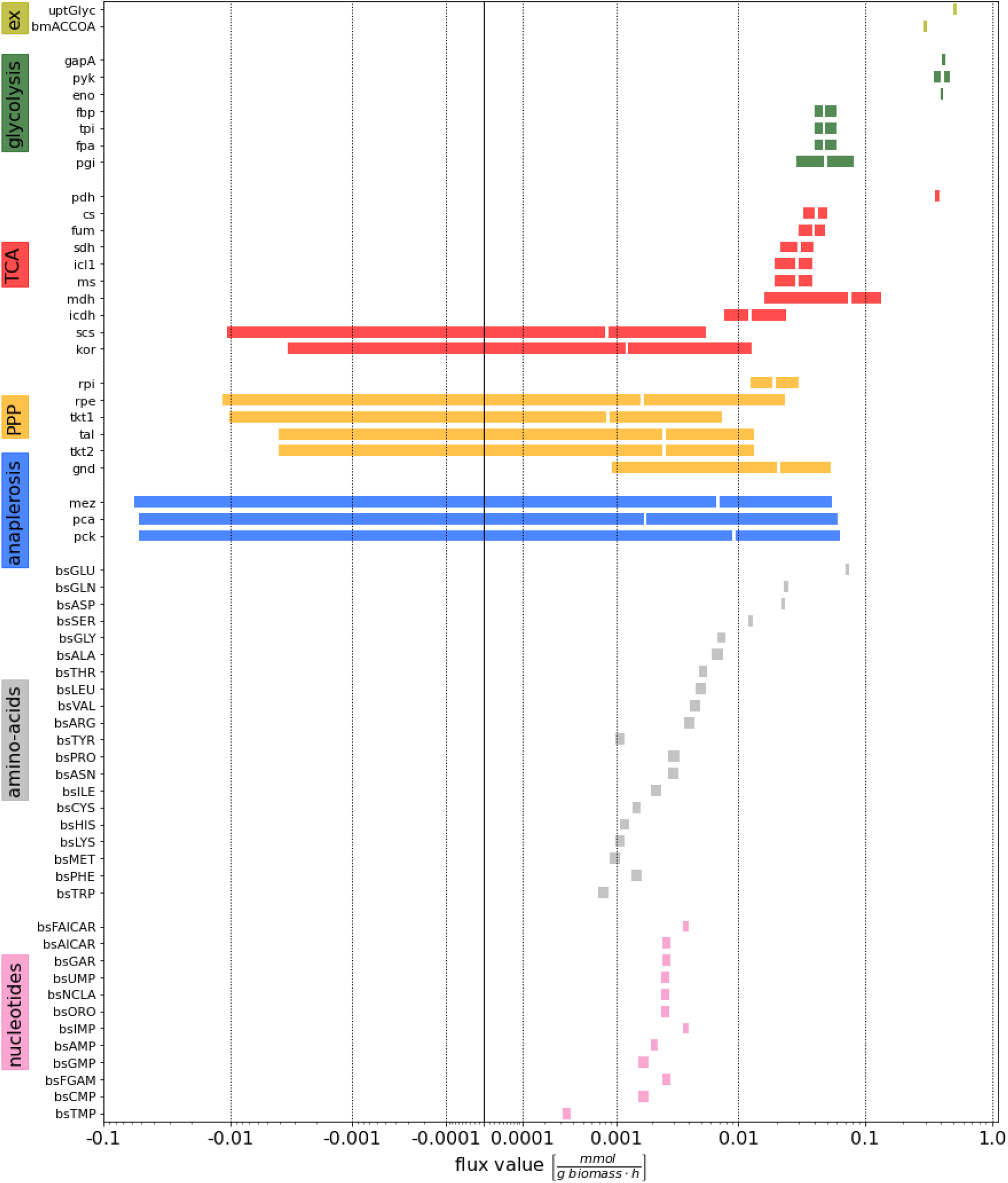
Credible intervals and expected values of absolute fluxes for BCG growing at 0.03 h^-1^ inferred with ^13^C^15^N-MFA. Colours indicate pathways and coloured bars specify 95% credible intervals (CrI) for net fluxes, with inscribed white line indicating the expected value (EV, in case of very narrow CrIs the EV is not displayed). The flux values are bi-symmetrically log transformed (*64*).

There are, however, also differences in the flux maps of central C metabolism. Our previous flux map showed cyclic fluxes around fructose 6-phosphate (F6P) node, involving the fluxes *pgi*, *gnd*, *tkt2*, *tal* and *tkt1*. This cycle is also confirmed by the present ^13^C^15^N-MFA study but with fluxes lower than previously reported. Notice, however, that the previously reported fluxes of the PPP represent best-fit values that were fixed in the statistical analysis due to their non-identifiability (22), and thus should serve for a qualitative comparison only. More accentuated differences are found in the fluxes of lower glycolysis. This is explained by the differences in the experimental setup between the two studies: In this study, tyloxapol was used as dispersant in the medium as replacement for tween-80 or oleic acid, which was a medium component in our previous study, and is known to be a C source for mycobacteria (*35*). Our results reveal that the lack of oleic acid as C source is compensated by an increase in lower glycolytic flux. It is interesting that whether using tween or tyloxapol in the medium there is no “global” effect on C fluxes under the investigated conditions.

As in our former ^13^C-MFA analysis (22), net fluxes of the anaplerotic reactions, pyruvate carboxylase (*pca*), PEP carboxykinase (*pck*), and malic enzyme (*mez*), could not be resolved as seen from their large CrIs in Fig. 4. This is a consequence of the cyclic network topology of the anaplerotic node (*36*). In our previous analysis *pca* and *mez* were aggregated by lumping oxaloacetate (OAA) and malate (MAL), but in this study the three anaplerotic reactions were modelled in detail without imposing any assumption on their (bi)directionalities. Despite limited information in the co-labelling data for identification of the directionalities of the anaplerotic reactions, we were able to improve the resolution of the flux map and limit the absolute flux values to a narrow range of ± 0.06 mmol g_biomass_^-1^ h^-1^. Furthermore, the two-dimensional (2D) marginal posterior probability distributions provide new insights into anaplerotic flux correlations (Fig. 5), which effectively narrows down the joint space of possible values to concise ring-like areas of the flux space. From our analyses many flux constellations can be ruled out. For instance, it is unlikely that two of the reactions, *pca* and *pck* or *pck* and *mez*, both carry zero flux. The flux pairs *pck*/*mez* and *pck*/*pca* are largely positively correlated, meaning that a larger value of one flux implies a larger value of the other. In both cases, two different possible modes emerge, explaining the data equally well, one with larger and one with smaller values of *mez* and *pca*, respectively. In contrast, *pca* and *mez* are largely negatively correlated. The results show that at least one of the three anaplerotic reactions is operating in gluconeogenetic direction. In a larger context, Fig. S4 shows that *mdh* and *pyk* are highly correlated with *mez* and *pck*, respectively. In conclusion, the C flux profile for BCG growing at a faster growth rate (0.03h^-1^) inferred here by BMA-based ^13^C^15^N-MFA represents an independent replication of the previous ^13^C-MFA-derived C fluxes. The comparable C flux maps derived from the two labelling approaches and the two MFA platforms validate the application of our Bayesian approach, which imposes fewer modeling assumptions, and delivers an increased flux resolution for the previously non-inferable anaplerotic cyclic nodes.

**Fig. 5.**
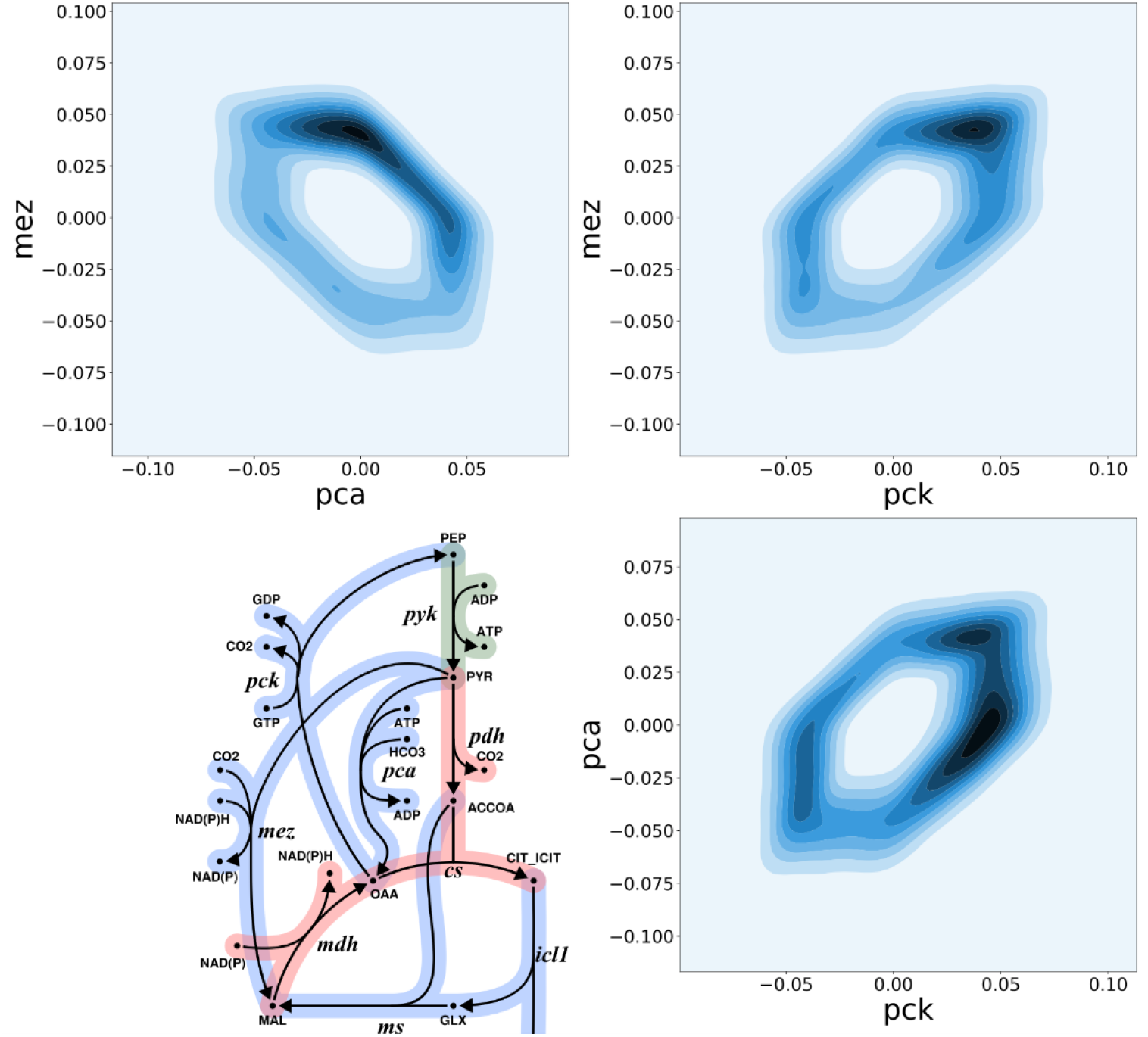
Joint marginal posterior probability plots of the fluxes of the three anaplerotic reactions pyruvate carboxylase (*pca*), PEP carboxykinase (*pck*), and malic enzyme (*mez*). The metabolic network of the anaplerotic node is shown in the lower left. Darker (lighter) colours indicate regions of higher (lower) probability according to the labelling data. For *mez* vs *pca*, *mez* vs *pck*, and *pca* vs *pck* flux correlations have ring-like shapes, giving raise to two modes of negative or positive correlation. See supplementary Fig. S4 for an extended version.

### ^13^C^15^N-MFA quantifies CN-fluxes, and thereby N-fluxes

In addition to the C fluxes discussed in the previous section, ^13^C^15^N-MFA together with the extended scope of the network allowed us, for the first time, to quantify CN net fluxes in *M. bovis* BCG, i.e. reaction fluxes in the amino acid and nucleotide biosynthetic pathways. These fluxes are shown in Fig. 3 (hexagons with thick red borders) and Fig. 4 in relative and absolute numbers. The largest CN flux is *bsGLU* (glutamate dehydrogenase), with an EV of 0.072 mmol g biomass^-1^ h^-1^ (95% CrI: 0.071 – 0.074 mmol g biomass^-1^ h^-1^), followed by *bsGLN* (glutamine synthetase), *bsASP* (aspartate transaminase) and *bsSER* (serine deaminase). The CN-fluxes of the remaining amino acid and nucleotide synthesis including adenosine monophosphate (AMP), guanosine monophosphate (GMP), cytidine monophosphate (CMP), inosine monophosphate (IMP) and uridine monophosphate (UMP), are one order of magnitude lower, with values rendering the proportion to which they contribute to biomass formation (Fig. 4).

When comparing CN fluxes with the C fluxes in Fig. 4, or with any previously reported flux values, it is important to realize that fluxes must be interpreted in relation to the actual material flows they represent. For example, it has been previously reported (*22*), that alanine aminotransferase (*bsALA*) is the largest biosynthetic net carbon flux, being one order of magnitude larger than glutamate dehydrogenase (*bsGLU*). In this study, we found that *bsALA* (95% CrI: 0.006 – 0.007 mmol g biomass^-1^ h^-1^) is one order of magnitude *lower* than *bsGLU*. This apparent discrepancy is explained by the nitrogen demand that is not within the scope of ^13^C-MFA. More precisely, conventional ^13^C-MFA can only give a lower bound for the nitrogen requirement of an amino acid by summing up the (C) fluxes of reactions that incorporate nitrogen from the donor. For example, adding the fluxes to alanine (ALA), asparagine (ASN), glutamate, glutamine (GLN), leucine (LEU), lysine (LYS), phenylalanine (PHE), serine (SER), tyrosine (TYR), tryptophan (TRP), valine (VAL), ornithine (ORN), arginine (ARG), and the contributions to nucleotide synthesis, gives a value of 0.061 mmol g biomass^-1^ h^-1^, which underestimates the EV of the *bsGLU* net CN flux 0.071 mmol g biomass^-1^ h^-1^ derived by ^13^C^15^N-MFA in this study.

The strength of ^13^C^15^N-MFA is that it provides quantitative flux measurements for N metabolism, via the CN fluxes. The N flux map shown in Fig. 6 highlights the central role of glutamate (GLU) as a N donor. To a lesser extent, glutamine (GLN) and aspartate (ASP) are also N donors, which explains their significantly higher CN flux as compared to the C fluxes previously reported (*22*). Glutamate donates its N to other amino acids through various transamination reactions. The centrality of this node for N assimilation was experimentally confirmed by examining substrate utilization of a glutamate auxotroph of *M. bovis* BCG with a transposon mutation in gltBD, a gene encoding glutamine oxoglutarate aminotransferase (GOGAT) that catalyzes the synthesis of GLU from OXG and GLN (*37*). Whereas the wild type *M. bovis* BCG strain could grow with GLYC as sole C and NH_3_, ASP, GLU and GLN as sole N sources (slope m >0), the gltBD mutant was able to grow only on glutamate as the N source (Fig. 7). The CN-fluxes for GLU and ASP-derived amino acids including ASN, threonine (THR), isoleucine (ILE), LYS, methionine (MET), ARG and PRO and phosphoglyceric acid (PGA)/phosphoenlypuruvate (PEP) derived ALA, SER, cysteine (CYS), VAL and leucine (LEU) are higher than the PPP-derived amino acids (Fig. 4). Interestingly, ALA had the largest pool size amongst the protein-derived amino acids, but the alanine aminotransferase CN flux was by far not the highest. Also, for the other amino acids, no direct relationship between pool sizes and fluxes is visible, highlighting that pool size and CN flux are complementary but independent measures of metabolism (Fig. 8; Fig. S5) (*38*).

**Fig. 6.**
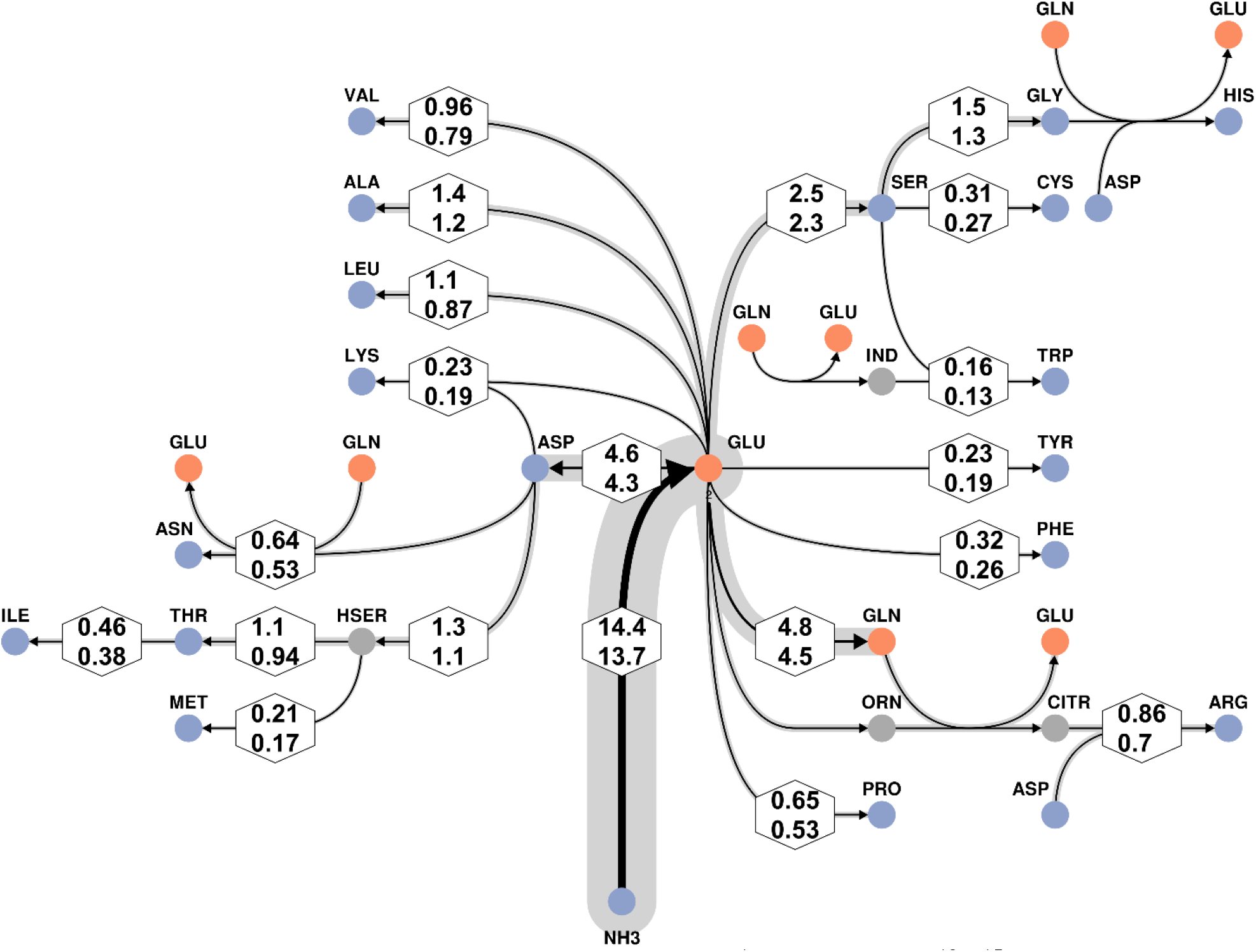
Bayesian N-flux map for BCG growing at 0.03 h^-1^ inferred with ^13^C^15^N-MFA. GLU-centric view with focus on amino acids. The line strengths code for the expected values (EV) of the net flux posterior probability distributions. Lower and upper limits of their associated 95% credible intervals (CrIs) are given in hexagons. To maintain compatibility with Fig. 3, values are given relative to the glycerol uptake flux (set to 100). Glutamate (GLU) is the primary nitrogen donor for most amino acids, but also glutamine (GLN), aspartate (ASP), and serine (SER) are important nitrogen hubs.

**Fig. 7.**
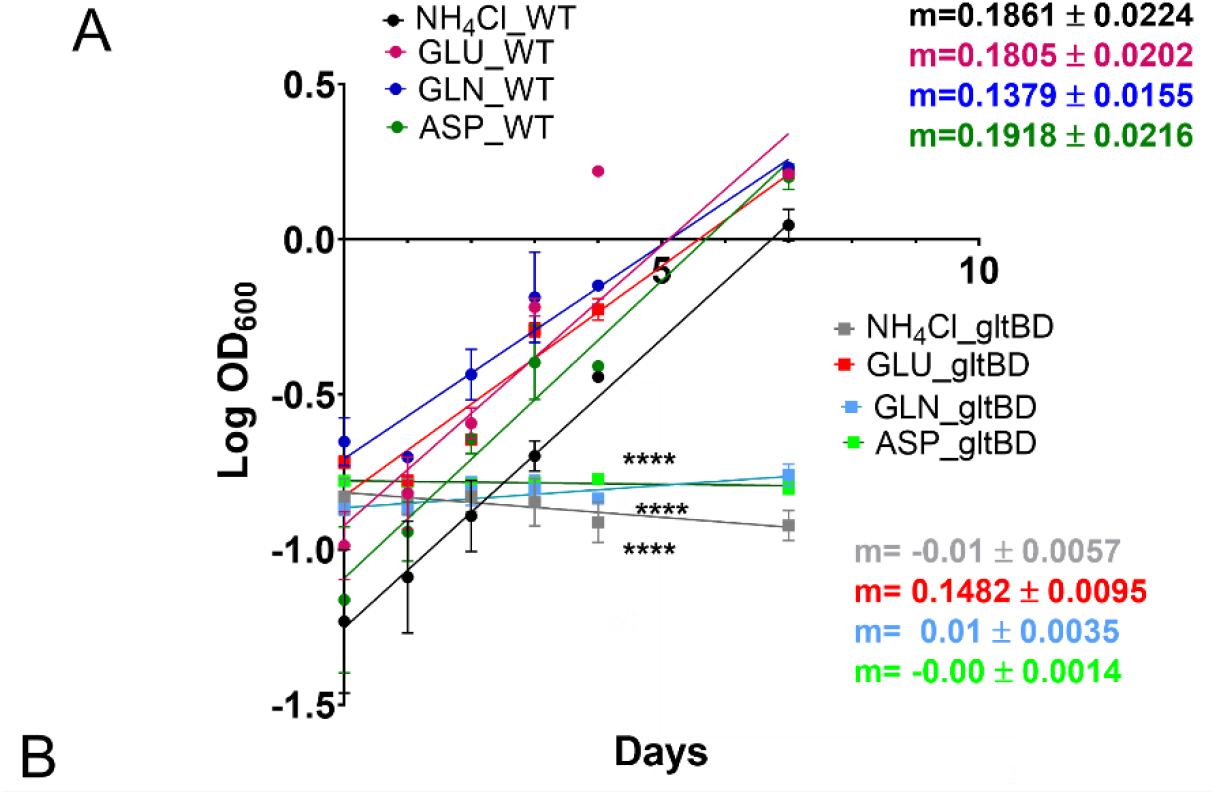
Growth of wild type and gltBD mutant BCG strain on minimal medium containing glycerol, GLU, ASP, GLN and NH_4_Cl. A positive slope m>0 indicates exponential growth. A negative slope (m<0) indicates no exponential growth. Values are mean ± SEM (n=3 independent replicates). * indicates statistically significant deviation of the slope from 0; ****, P<0.00001.

**Fig. 8.**
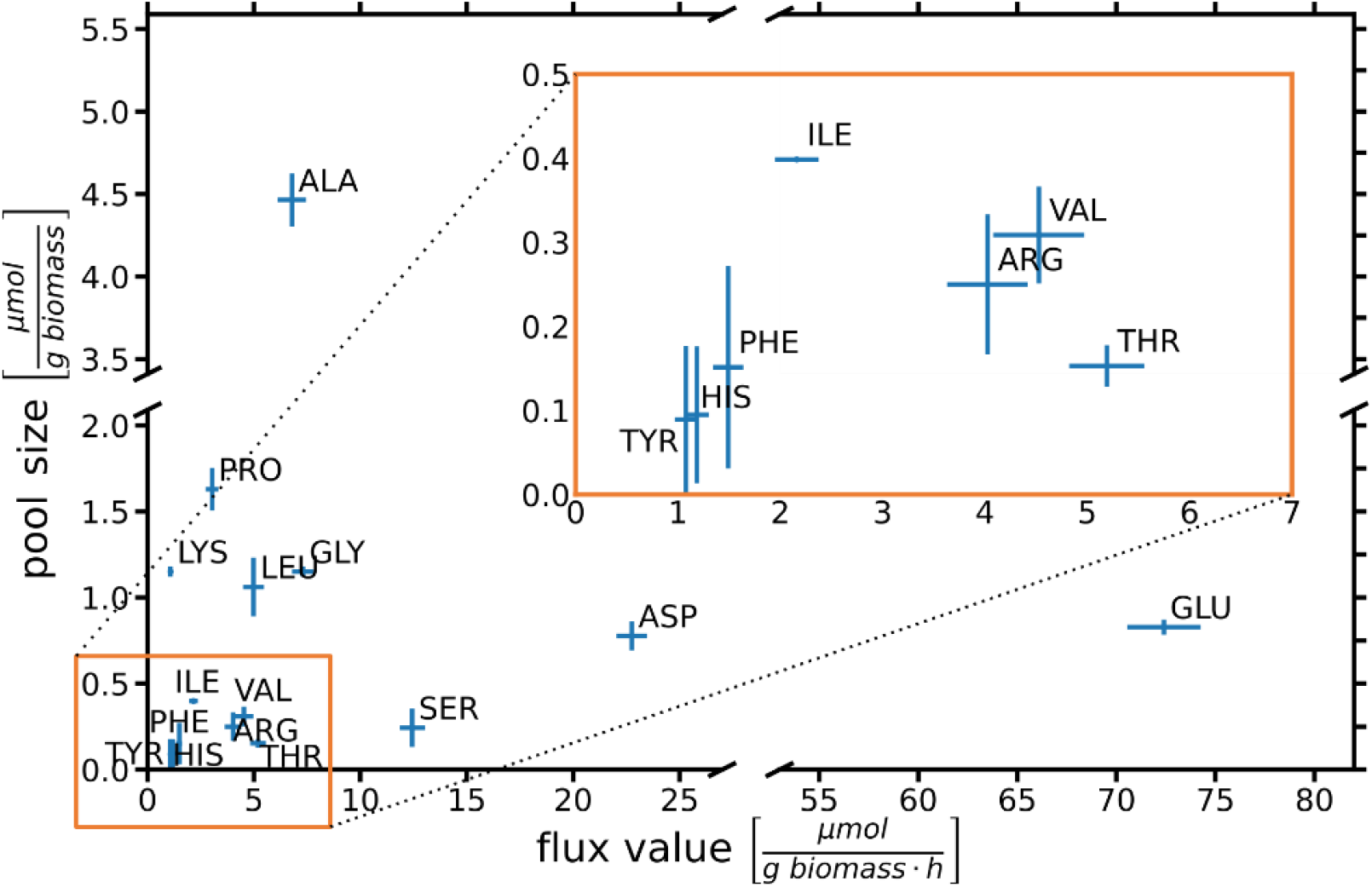
Pool sizes vs. biosynthetic CN fluxes of protein-derived amino acids. Values are mean ± 2 SD (n=3 independent measurements) for pool size measurements and 95% CrIs for flux values.

### Bayesian Multi-Model ^13^C^15^N-MFA uncovers reversibility of glycine biosynthesis and unidirectionality of leucine and isoleucine biosynthesis

GLU and GLN serve as the main amino donors for the synthesis of other amino acids in BCG (Fig. 4 and Fig. 6). To meet the demands for protein, RNA and DNA synthesis, the N net flux is principally directed towards amination. However, the reversibility of the enzymes responsible for catalyzing transaminases provide cells with the ability to adapt rapidly to environmental conditions such as changes in N availability. Indeed, fine-tuning the activities of transaminases to modulate C flux has been demonstrated in the CHO eukaryotic cell line (*39*). This means that while the biosynthetic net reaction flux is directed towards amination, a simultaneous forward and backward (bidirectional) flux is likely occurring *in vivo*.

Due to the lack of evidence about the reversibility of mycobacterial transaminases, all transamination reactions were modelled as bidirectional reaction steps, resulting in a flux estimation problem with 86 degrees of freedom (71 fluxes and 15 measurement group scales) to be recovered from 144 measurements (109 univariate MIDs and 34 rate and biomass efflux measurements). Here, BMA was used to minimise overfitting (see Materials and Methods). This technique can explore the space of all possible combinations of uni- and bidirectional reactions (each combination codes for a model) and weights each model variant by its ability to explain the labelling data (see also Fig. 2). In contrast to the conventionally used single-model approaches, the BMA-based ^13^C^15^N-MFA enables the rigorous statistical inference of reaction bidirectionality (*26*). The univariate MIDs did not allow all bidirectional reactions of biosynthesis to be classified as either bi- or unidirectional (Fig. S6) with distinct exceptions: glycine hydroxymethyltransferase (*bsGLY*) that catalyzes glycine (GLY) biosynthesis was determined to be bidirectional with 100 ± 0% probability. In contrast serine deaminase, isoleucine, and leucine transaminase (*bsSER*, *bsILE* and *bsLEU*) were found to be unidirectional with high probability (98.6 ± 0.3% probability, 99.8 ± 0.05%, and 100 ± 0% probability, respectively).

## Discussion

The interplay between C and N assimilation and dissimilation is required to sustain cell metabolism and function. To date, the knowledge about C and N co-assimilation and these fluxes within a cell remains limited. For measuring metabolic fluxes, isotope tracing studies in combination with computational modelling is a gold standard. Recently, multiple stable isotopic tracers have been used to measure C and N enrichments and to derive insights into N assimilation in eukaryotic systems including yeast, plant, and human cancer cells (*27*, *31*). Two pioneering studies determined metabolic fluxes of the GS-GOGAT pathway in *C. glutamicum* by INST ^15^N-MFA (*12*) and the CN fluxes for arginine metabolism in *K. lactis (15*). Both these studies had limitations; the former was limited to measuring CN fluxes in a small sub-network, and the latter ^13^C^15^N-flux analysis relied on a tailored labelling strategy that exploited the fact that the ^13^C tracer was incorporated faster into metabolites than the ^15^N tracer.

Here we generalized the well-established methodology of ^13^C-MFA towards ^13^C^15^N-MFA for deriving a system-wide CN, and thereby N fluxes from a combination of isotopic co-labelling experiments and comprehensive C and N atom transition modelling. The generalization is paired with the use of Bayesian model averaging, which provided rigorous quantification of CN metabolic fluxes with low- and medium-resolution MS data by measuring intracellular metabolic fluxes through the central C and N network of *M. bovis* BCG. Key advantages of BMA-based ^13^C^15^N-MFA are the extension of scope and refinement in terms of the metabolic network, while, simultaneously, tackling the consequent introduced uncertainties in the network formulation. This enabled the rigorous statistical assessment of all reactions, even those with unknown bidirectionality and cyclic network structures that were previously unresolvable.

We and others have demonstrated that the TB pathogen co-metabolizes multiple host nutrients during infection (*19*–*21*, *40*, *41*). Co-catabolism of C sources by mycobacteria and the associated metabolic regulations have been previously demonstrated by multiple studies using transcriptional, metabolomics and genetic approaches (*21*, *42*–*47*). Although multiple studies have explored the C and N metabolism of Mtb *in vitro* and within *ex vivo* and *in vivo* animal models, the information about CN and N fluxes and the key metabolic steps that could be targeted for drug development remain limited. Here we used [^13^C_3_]-glycerol and [^15^N_1_]-ammonium chloride dual isotopic labelling of steady state BCG cultures to measure intracellular CN and N fluxes. We provide the first comprehensive intracellular CN and N flux distributions in a biological system. A comparison with our previously published ^13^C-MFA in Mtb and *M. bovis* BCG (*22*) showed a broad agreement for the net fluxes of central C metabolism. C fluxes through glycolysis and PPP including phosphoglucose isomerase (*pgi*), fructose bisphosphatase (*fbp*), aldolase (*fba*), glucose-6-phosphate dehydrogenase/glucolactonase (*gnd*), transketolase (*tkt1*, *tkt2*) and transaldolase (*tal*) were not quantified accurately with our previous ^13^C-MFA and recent ^13^C^15^N-MFA study because PPP reactions lack of sufficient labelling information from proteinogenic amino acids. Incorporating labelling information from glycogen, ribose moiety of DNA, glucosamine moiety from peptidoglycan and lipopolysaccharides, such as suggested by (*48*) is a promising approach to improve flux precision in this area of metabolism. The fluxes of the three anaplerotic enzymes pyruvate carboxylase (*pca*), PEP carboxy kinase (*pck*), and malic enzyme (*mez*), which play an important role in metabolism connecting catabolism and anabolism with energy generation (*49*), are notoriously difficult to determine. We demonstrate that BMA-based ^13^C^15^N-MFA provides an effective tool to constrain the anaplerotic net fluxes. The Bayesian flux map is consistent with our previous results showing non-identifiability of individual anaplerotic fluxes (*22*), however, the co-labeling data in this study informs about distinct likely flux couplings of *pca*, *pck* and *mez* which could not be measured in our previous work. From our analysis we conclude that at least one of the three reactions is operating in the gluconeogenetic direction, and it is unlikely that two of the associated fluxes are zero at the same time.

Beyond C fluxes, we measured CN and N fluxes to amino acid and nucleotide (purine and pyrimidine) biosynthesis, providing novel information which cannot be deduced from our former ^13^C-MFA. We previously identified ASP, GLU and GLN as primary C and N sources for Mtb in human host macrophages (*19*, *20*). Here we quantified the CN and N flux for the biosynthesis of these amino acids. We identified glutamate biosynthesis *bsGLU* as the primary node for CN flux. This is consistent with the finding that ASP, GLN and NH_4_^+^ as sole N sources in Roisin’s minimal medium failed to rescue the growth of a *M. bovis* BCG mutant lacking functional *glt*BD gene and thereby lacking *de novo* glutamate synthesis (*37*, *50*). GLU is a well-established N source for *in vitro* and intracellular growth of mycobacteria. GLU metabolism is also crucial in mycobacteria to resist acidic and nitric oxide stress inside macrophages (*37*, *50*) and is therefore a prime metabolic and regulatory node. Furthermore, we were able to quantify bidirectionality probabilities for glycine hydroxymethyltransferase (*bsGLY*), serine deaminase (*bsSER*) and isoleucine and leucine transaminase (*bsILE* and *bsLEU*). C and N metabolic profiling has been attempted using isotopic labelling by other studies. Blank *et al*. investigated simultaneous C and N incorporation in *Saccharomyces cerevisiae* administering two different isotopic substrates ^13^C-glucose and ^15^N-alanine and measured dual label incorporation in amino acids using FT-ICR-MS (*31*). Our study demonstrates that conventional GC-MS, which is the traditional workhorse for isotopomer analysis, can also be used to derive dual labelled data sets and univariate MIDs of amino acids for robust and reproducible flux quantification by ^13^C^15^N-MFA. Our CN metabolic network is currently limited to amino acid and nucleotide biosynthesis, but there are further scopes for extension of this model through addition of biocomponents such as co-factors NADH and NADPH, and lipids that requires CN assimilation and dissimilation.

In summary, we have developed Bayesian ^13^C^15^N-MFA, a powerful tool for simultaneous quantification of intracellular C, CN and N metabolic fluxes in a living system. We applied it to M. *bovis* BCG. ^13^C^15^N-MFA identified glutamate as the central CN node, revealed the most likely operational modes of the anaplerotic fluxes, and resolved the uni/bidirectionalities of glycine, serine, isoleucine, and leucine biosynthesis. Our ^13^C^15^N-MFA workflow described here is applicable to any C and N isotopic co-labelling experiment, and the computational platform developed in this work allows analyses of low- and medium-resolution MS data to provide rigorous quantification of CN metabolic fluxes in any biological system.

## Acknowledgements

The authors are thankful to Wolfgang Wiechert for excellent working conditions at the IBG-1. We are grateful to VIB - Vlaams Instituut voor Biotechnologie for providing us *M. bovis* BCG gltBD transposon mutant (BCG_3922c TnInsertion-8654).

## Author contributions

Conceptualization: JM, KN. Experimental analyses, and data collection: KB, YX, JB, CC, JN. Data analyses and visualization: MB, KB. Computational analyses: MB, KB. Implementation of computational framework: MB, AT. Writing original draft: KB, JM, KN, MB. Writing review and editing: KB, MB, CC, YX, JN, JB, DB, MJB, KN, JM. Funding acquisition: MJB, JM.

## Competing interests

The authors declare no competing interests exist.

## Funding

This study was supported by Biotechnology and Biological Sciences Research Council (BBSRC) grants BB/L022869/1 and BB/V010611/1 and the Electronics and Physical Sciences Research Council grants EP/R031118/1 and EP/P001440/1, United Kingdom.

## Data availability

Mass spectrometry data and ^13^C-^15^N atom transition model is provided as supplementary material.

## Materials and Methods

### Bacterial strains

*Mycobacterium bovis* BCG Pasteur, which was originally purchased from the American Type Culture Collection (ATCC 35748), was used for this study. *M. bovis* BCG gltBD transposon mutant (BCG_3922c TnInsertion-8654) was procured from VIB - Vlaams Instituut voor Biotechnologie.

### Media composition

Middlebrook 7H11 agar and Middlebrook 7H9 broth containing 5% (vol/vol) oleic acid-albumin-dextrose-catalase enrichment medium supplement (Becton Dickenson) and 0.5% (vol/vol) glycerol were used to grow cultures from frozen stocks and for counting the numbers of culturable bacteria in chemostat samples. Brain heart infusion agar was used to assess culture purity (Sigma Aldrich). For cultivation of *M. bovis* BCG in the bioreactor, roisins minimal medium with composition-KH_2_PO_4_, 1 g litre^-1^; Na_2_HPO_4_, 2.5 g litre^-1^; NH_4_Cl, 5.9 g litre^-1^; K_2_SO_4_, 2 g litre^-1^; ZnCl_2_, 0.08 mg litre^-1^; FeCl_3_, 0.4 mg litre^-1^; CuCl_2_, 0.02 mg litre^-1^; MnCl_2_, 0.02 mg litre^-1^; Na_2_B4O_7_, 0.02 mg litre^-1^; NH_4_MoO_4_, 0.02 mg litre^-1^; MgCl_2_, 0.0476 g litre^-1^; CaCl_2_, 0.055 g litre^-1^; Tyloxapol, 01% (v/v); Glycerol, 0.5% (v/v).

### Growth of *M. bovis* in the bioreactor and chemostat

*M. bovis* BCG strain was cultured in a 2 litre bioreactor (Electrolab Fermac 310/60) under growth conditions (Table S1). Cultures were grown as batch for 7 days. Continuous cultures were grown under chemostat conditions at a growth rate of 0.03 h^-1^ maintained by the media flow rate (*22*). Media was pumped into the chemostat using a peristaltic pump (Rainin Rabbit Plus). Cultures were grown for 3-4 volume changes in the unlabelled media to assure a metabolic steady-state before introducing isotopically labelled media. [^13^C_3_] glycerol (12.5%) (Purchased from CK Isotopes, 99% purity) and [^15^N_1_] NH_4_Cl (20%) (Purchased from Merck, 98% atom purity) were the carbon and nitrogen isotopically labelled substrates in the media. Isotopic stationary state was assessed by measuring % label in the proteinogenic amino acids of cultures drawn at different times during label feed (Fig. S1C).

### Chemostat measurements and culture analyses

Cultures were monitored every day to check for contamination by plating on BHI agar media and ziehl neelsen stain. Cultures from chemostat were regularly sampled for measuring OD (spectrophotomter from Thermo scientific) and colony forming units (*22*). Carbon-di-oxide production from the cultures was monitored using Gas analyser (Electrolab Fermac 368). The supernatant was collected, filtered using 0.22 μ unit filters and used for substrate consumption and product excretion analyses. To measure the glycerol uptake, the amounts of glycerol in the supernatant and fresh medium was measured using glycerol assay kit (Sigma Aldrich) by a coupled enzyme assay involving glycerol kinase and glycerol phosphate oxidase, resulting in a colorimetric (570 nm) product, proportional to the glycerol present. To measure NH_4_Cl uptake, the amounts of NH_4_Cl was measured using ammonia assay kit (Sigma Aldrich) by reaction of ammonia present in the samples involving L-glutamate dehydrogenase activity. Dry weight of the cells was measured by centrifuging cultures, drying the cell pellet using freeze dryer and weighing the cells. The dried pellet was used for protein analysis using Bicinchoninic Acid Kit for Protein Determination (Sigma Aldrich).

### Metabolite extraction

Labelled chemostat cultures were quenched using methanol:chloroform:water (2:1:2) extraction. Briefly, cultures were filtered using membrane Filter, 0.22 μm pore size, filter apparatus (Merck) and the filter was immersed into methanol:chloroform, mixed, incubated on ice for 30 minutes and water was added, followed by centrifuging at room temperature for triphasic metabolite separation. The upper phase was collected separately and dried and used for mass spectrometry analysis. The lower and intermediate phase were mixed into one phase by addition of one more volume of methanol and chloroform and centrifuged for 30 minutes at room temperature. Supernatant was discarded, the pellet was hydrolysed in 6 M hydrochloric acid for 24 hours at 100°C and the hydrolysate was dried for mass spectrometry analysis.

### Mass spectrometry analysis of amino acids

Dried upper phase was derivatised using N-Methyl-N-(trimethylsilyl)trifluoroacetamide, MSTFA (sigma Aldrich) and dried hydrolysate were derivatised using tert-Butyldimethylsilyl chloride (TBDMSCl) were analysed using GC-MS (7890-5795 system) (*20*) Mass spectra were baseline corrected using MetAlign and mas isotopomer distribution (MID) data were extracted using the chemstation software. Identification of metabolites was done using NIST databases, literatures, and qualifier masses. Average ^13^C^15^N fractional abundances were calculated from two independent chemostat cultivations (three- or four technical replicates each) and quantitation of metabolite pool sizes was done using calibration curves (*20*). Further confirmation of ^13^C and ^15^N labelling in the amino acids were done using LC-MS orbitrap (Fig. S7). Briefly, hydrophilic interaction liquid chromatography (HILIC) was carried out on a Dionex UltiMate 3000 RSLC system (Thermo Fisher Scientific, Hemel Hempstead, UK) using a C18 and ZIC-pHILIC column (150 mm × 4.6 mm, 5 μm column, Merck Sequant). The column was maintained at 30°C and samples were eluted with a linear gradient (20 mM ammonium carbonate in water, A and acetonitrile, B) over 26 min at a flow rate of 0.3 ml/min. The injection volume was 10 μl and samples were maintained at 4°C prior to injection. For the MS analysis, a Thermo Orbitrap Q Exactive Plus (Thermo Fisher Scientific) was operated in polarity switching mode and the MS settings were used with resolution 70,000, AGC 106, m/z range 70–1400, sheath gas 40, Auxiliary gas 5, sweep gas 1, probe temperature 150°C and capillary temperature 275°C. For positive mode ionisation: source voltage +4.5 kV, capillary voltage +50 V, tube voltage +70 kV, skimmer voltage +20 V. For negative mode ionisation: source voltage-3.5 kV, capillary voltage-50 V, tube voltage-70 V, skimmer voltage-20 V. The data shown in supplementary Fig. S7 is a mass spectrum showing the multivariate ^13^C and ^15^N species identification for alanine.

### ^13^C^15^N-Metabolic flux analysis

Metabolic network model: The metabolic model *M. bovis* BCG used for the analysis was constructed using the network editor Omix v.2.0.7 (Omix Visualization GmbH & Co. KG, Lennestadt/Germany) (*51*), according to the protocol described in (*52*), based on the GSMN-TB genome-scale model of *M. tuberculosis* (*53*). The constructed model includes reactions of glycolysis, the PPP, the TCA cycle, anaplerosis, nucleotide and amino acid biosynthesis. Uptake reactions were considered for GLYC and ammonium chloride (NH_4_Cl). All biosynthesis pathway fluxes relevant for growth of *M. bovis* are modelled as effluxes, whose values represent their share in the biomass composition (*54*). Reactions were considered bidirectional including all transaminases, unless evidence was found that the reactions operate far from thermodynamic equilibrium under *in vivo* conditions (supplementary Table S2). Only in the latter case were those reactions modelled unidirectional. Each metabolic reaction was supplemented with carbon and nitrogen atom transitions, following the InChI atom enumeration scheme (*55*). Carbon symmetries of succinate (SUCC), fumarate (FUM), and diaminopimelic acid (DAP) were accounted for by the formulation appropriate label scrambling reactions. In total, the *M. bovis* BCG model consists of 248 metabolites (121 balanced intracellular and 127 unbalanced extracellular pools) and 184 metabolic reactions (149 unidirectional, 35 bidirectional). The most comprehensive model in the model set has 71 independent flux parameters (36 net and 35 exchange fluxes). The corresponding CN atom transition network model is formulated in the standardized document format for isotope-based MFA, FluxML v3 (*56*), and is found in supplementary Data S1.

Measurement models: In total 30 biomass effluxes were considered as measurements that were either obtained from biomass hydrolysates or calculated from intermediates (*22*) and supplied with Gaussian error of 5%. Uptake rates for glycerol and NH4Cl were fixed for the analysis. Labelling measurements of 15 amino acids, were corrected for the effect of natural abundant isotopes (*57*), adding up to 109 univariate MIDs (supplementary Table S3). The associated measurement errors were derived from up to 8 replicates (two independent chemostats, each with two-four replicate measurements). The complete measurement specification is given in the FluxML model file in supplementary Data S1.

Labelling system & simulation: The ^13^C-MFA high-performance simulator 13CFLUX2 v2.2 (*58*) was extended to simulate ^13^C^15^N isotopologues. Briefly, the essential cumomer framework (*59*) was generalized from single-atom to multi-atom species labelling systems. The resulting balance equations consist of a set of sparse linear equation systems that were solved sequentially with on-the-fly algebraic simplification guaranteeing numerical stability, accuracy, and efficiency. The resulting reduced co-labelling system has a state-space dimension of 471 (a reduction of a factor of 2,563 compared to the full co-labelling system).

Flux inference with Bayesian Model Averaging: Instead of conventional optimization-based single-model flux inference, in this work metabolic fluxes were estimated using a Bayesian multi-model approach. More precisely, net fluxes and reaction bidirectionalities were inferred simultaneously by employing BMA, implemented using a tailored MCMC approach (*26*). Herein, 13CFLUX2 was used for likelihood computation. For speed, sampling algorithms implemented in the C++ library HOPS v2.0.0 (*60*) were employed after suitable preprocessing using Polyround v0.1.8 (*61*). Parallel tempering with dynamic temperature selection was applied to sample from potentially multi-modal distributions. In total, 10 parallel chains were run from independent starting points, where per chain 15×10^6^ forward simulations were performed. For each chain, the first 5×10^6^ samples were discarded (burn-in). Proper convergence of the MCMC runs was diagnosed by measuring the Potential Scale Reduction Factor (PSRF) (*62*) on a subset of samples, where all but every 2,000th sample was disregarded (thinning). Computations were run on a workstation with dual Intel(R) Xeon(R) Gold CPU (61300 @ 2.8 GHz). PSRF, mixing plots in parameter as well as model spaces are provided in supplementary Table S4, Fig. S8 and Fig S9.

Statistical evaluation: From the MCMC results posterior probability distributions *p*(*v*|*D*) for the net fluxes *v* were derived, according to (EQ 1). Formally, the posterior probability of a model *M_i_* out of the model set {*M_i_*|*i* = 1,…, *N*} is given by

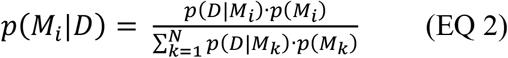

where *p*(*M_i_*), the prior knowledge about model *M_i_*, is considered equal for all models. *p*(*D*|*M_i_*) represents the high dimensional marginalization integral over all possible fluxes the model *M_i_* can take. The 95% CrIs were determined for each net flux (i.e. the range that contains the flux with a probability of 95% in view of the data), discarding the upper and lower 2.5% of the values. In addition, as point estimator, the expected value (EV) for the net flux is reported. The marginal distributions for the net fluxes are provided in supplementary Fig. S2. For each reversible reaction, the posterior probability of the reaction being bidirectional was determined, according to (EQ 2), as ratio of models sampled with the reaction set bidirectional divided by the total number of sampled models (supplementary Fig. S6).

### Statistical Analysis

Students-t-test, analysis of variance (ANOVA) and linear regression analyses was performed in Graphpad Prism 8.0.

## Notes

### Competing Interest Statement

The authors have declared no competing interest.

## References

1. S. Niedenführ, W. Wiechert, K. Nöh, How to measure metabolic fluxes: A taxonomic guide for 13C fluxomics. Current Opinion in Biotechnology. 34 (2015), pp. 82–90.

2. J. Nielsen, It Is All about Metabolic Fluxes. Journal of Bacteriology. 185 (2003), pp. 7031–7035.

3. W. Wiechert, Metabolic Flux Analysis. Metab. Eng. 3(3):195–206. doi: 10.1006/mben.2001.0187. (2001).

4. N. Zamboni, S.-M. Fendt, M. Rühl, U. Sauer, 13C-based metabolic flux analysis. Nature Protocols. 4 (2009), doi:10.1038/nprot.2009.58.

5. K. Sonntag, L. Eggeling, A. A. de Graaf, “Flux partitioning in the split pathway of lysine synthesis in Corynebacterium glutamicum Quantification by 13C-and ‘H-NMR spectroscopy” (1993).

6. W. Wiechert, A. A. de Graaf, “Bidirectional Reaction Steps in Metabolic Networks: I. Modeling and Simulation of Carbon Isotope Labeling Experiments” (1997).

7. V. Chubukov, L. Gerosa, K. Kochanowski, U. Sauer, Coordination of microbial metabolism. Nature Reviews Microbiology. 12 (2014), pp. 327–340.

8. M. P. Tan, P. Sequeira, W. W. Lin, W. Y. Phong, P. Cliff, S. H. Ng, B. H. Lee, L. Camacho, D. Schnappinger, S. Ehrt, T. Dick, K. Pethe, S. Alonso, Nitrate respiration protects hypoxic Mycobacterium tuberculosisagainst acid- and reactive nitrogen species stresses. PLoS ONE. 5 (2010), doi:10.1371/journal.pone.0013356.

9. K. Kurmi, M. C. Haigis, Nitrogen Metabolism in Cancer and Immunity. Trends in Cell Biology. 30 (2020), pp. 408–424.

10. W. Wiechert, K. Nöh, Isotopically non-stationary metabolic flux analysis: Complex yet highly informative. Current Opinion in Biotechnology. 24 (2013), pp. 979–986.

11. K. Nöh, W. Wiechert, The benefits of being transient: Isotope-based metabolic flux analysis at the short time scale. Applied Microbiology and Biotechnology. 91 (2011), pp. 1247–1265.

12. M. Tesch, A. A. de Graaf, H. Sahm, “In Vivo Fluxes in the Ammonium-Assimilatory Pathways in Corynebacterium glutamicum Studied by 15 N Nuclear Magnetic Resonance” (1999).

13. P. Goel, M. Bhuria, M. Kaushal, A. K. Singh, Carbon: Nitrogen interaction regulates expression of genes involved in N-Uptake and assimilation in brassica juncea L. PLoS ONE. 11 (2016), doi:10.1371/journal.pone.0163061.

14. M. R. Naliwajski, M. Skłodowska, The relationship between carbon and nitrogen metabolism in cucumber leaves acclimated to salt stress. PeerJ. 2018 (2018), doi:10.7717/peerj.6043.

15. G. Romagnoli, M. D. Verhoeven, R. Mans, Y. Fleury Rey, R. Bel-Rhlid, M. van den Broek, R. Maleki Seifar, A. ten Pierick, M. Thompson, V. Müller, S. A. Wahl, J. T. Pronk, J. M. Daran, An alternative, arginasein-dependent pathway for arginine metabolism in Kluyveromyces lactis involves guanidinobutyrase as a key enzyme. Molecular Microbiology. 93, 369–389 (2014).

16. World Health Organization., Global tuberculosis report 2020. (World Health Organization, 2020).

17. J. C. Palomino, A. Martin, Drug resistance mechanisms in Mycobacterium tuberculosis. Antibiotics. 3 (2014), pp. 317–340.

18. S. G. Kurz, J. J. Furin, C. M. Bark, Drug-Resistant Tuberculosis: Challenges and Progress. Infectious Disease Clinics of North America. 30 (2016), pp. 509–522.

19. D. J. V. Beste, K. Nöh, S. Niedenführ, T. A. Mendum, N. D. Hawkins, J. L. Ward, M. H. Beale, W. Wiechert, J. McFadden, 13C-flux spectral analysis of host-pathogen metabolism reveals a mixed diet for intracellular mycobacterium tuberculosis. Chemistry and Biology. 20, 1012–1021 (2013).

20. K. Borah, M. Beyß, A. Theorell, H. Wu, P. Basu, T. A. Mendum, K. Nöh, D. J. V. Beste, J. McFadden, Intracellular Mycobacterium tuberculosis Exploits Multiple Host Nitrogen Sources during Growth in Human Macrophages. Cell Reports. 29, 3580–3591.e4 (2019).

21. L. P. S. de Carvalho, S. M. Fischer, J. Marrero, C. Nathan, S. Ehrt, K. Y. Rhee, Metabolomics of mycobacterium tuberculosis reveals compartmentalized co-catabolism of carbon substrates. Chemistry and Biology. 17, 1122–1131 (2010).

22. D. J. V. Beste, B. Bonde, N. Hawkins, J. L. Ward, M. H. Beale, S. Noack, K. Nöh, N. J. Kruger, R. G. Ratcliffe, J. McFadden, 13c metabolic flux analysis identifies an unusual route for pyruvate dissimilation in mycobacteria which requires isocitrate lyase and carbon dioxide fixation. PLoS Pathogens. 7 (2011), doi:10.1371/journal.ppat.1002091.

23. K. Borah, T. A. Mendum, N. D. Hawkins, J. L. Ward, M. H. Beale, G. Larrouy-Maumus, A. Bhatt, M. Moulin, M. Haertlein, G. Strohmeier, H. Pichler, V. T. Forsyth, S. Noack, C. W. Goulding, J. McFadden, D. J. v Beste, Metabolic fluxes for nutritional flexibility of Mycobacterium tuberculosis. Molecular Systems Biology. 17 (2021), doi:10.15252/msb.202110280.

24. N. Zamboni, S.-M. Fendt, M. Rühl, U. Sauer, 13C-based metabolic flux analysis. Nature Protocols. 4 (2009), doi:10.1038/nprot.2009.58.

25. C. P. Long, M. R. Antoniewicz, High-resolution 13C metabolic flux analysis. Nature Protocols. 14 (2019), doi:10.1038/s41596-019-0204-0.

26. A. Theorell, K. Nöh, Reversible jump MCMC for multi-model inference in Metabolic Flux Analysis. Bioinformatics. 36, 232–240 (2020).

27. R. Nilsson, M. Jain, Simultaneous tracing of carbon and nitrogen isotopes in human cells. Molecular BioSystems. 12, 1929–1937 (2016).

28. M. I. Borkum, P. N. Reardon, R. C. Taylor, N. G. Isern, Modeling framework for isotopic labeling of heteronuclear moieties. Journal of Cheminformatics. 9 (2017), doi:10.1186/s13321-017-0201-7.

29. J. Kappelmann, M. Beyß, K. Nöh, S. Noack, Separation of 13C- And 15N-Isotopologues of Amino Acids with a Primary Amine without Mass Resolution by Means of O-Phthalaldehyde Derivatization and Collision Induced Dissociation. Analytical Chemistry. 91, 13407–13417 (2019).

30. J. Kappelmann, B. Klein, P. Geilenkirchen, S. Noack, Comprehensive and accurate tracking of carbon origin of LC-tandem mass spectrometry collisional fragments for 13C-MFA. Analytical and Bioanalytical Chemistry. 409, 2309–2326 (2017).

31. L. M. Blank, R. R. Desphande, A. Schmid, H. Hayen, Analysis of carbon and nitrogen co-metabolism in yeast by ultrahigh-resolution mass spectrometry applying 13C- and 15N-labeled substrates simultaneously. Analytical and Bioanalytical Chemistry. 403, 2291–2305 (2012).

32. T. Grotkjær, M. Åkesson, B. Christensen, A. K. Gombert, J. Nielsen, Impact of Transamination Reactions and Protein Turnover on Labeling Dynamics in 13C-Labeling Experiments. Biotechnology and Bioengineering. 86, 209–216 (2004).

33. W. Wiechert, The thermodynamic meaning of metabolic exchange fluxes. Biophysical Journal. 93, 2255–2264 (2007).

34. J.A. Hoeting, D. Madigan, A.E. Raftery, C.T. Volinsky. Bayesian model averaging: A tutorial. Stat. Sci. 14, 382–417 (1999).

35. R. D. Pietersen, I. du Preez, D. T. Loots, M. van Reenen, D. Beukes, G. Leisching, B. Baker, Tween 80 induces a carbon flux rerouting in Mycobacterium tuberculosis. Journal of Microbiological Methods. 170 (2020), doi:10.1016/j.mimet.2019.105795.

36. J. Kappelmann, W. Wiechert, S. Noack, Cutting the Gordian Knot: Identifiability of Anaplerotic Reactions in Corynebacterium glutamicum by Means of 13 C-Metabolic Flux Analysis. Biotechnol. Bioeng. 113, 661–674 (2016).

37. A. J. Viljoen, C. J. Kirsten, B. Baker, P. D. van Helden, I. J. F. Wiid, The role of glutamine oxoglutarate aminotransferase and glutamate dehydrogenase in nitrogen metabolism in Mycobacterium bovis BCG. PLoS ONE. 8 (2013), doi:10.1371/journal.pone.0084452.

38. Y. Wang, F. E. Wondisford, C. Song, T. Zhang, X. Su, Metabolic flux analysis—linking isotope labeling and metabolic fluxes. Metabolites. 10 (2020), pp. 1–21.

39. J. Wahrheit, J. Niklas, E. Heinzle, Metabolic control at the cytosol-mitochondria interface in different growth phases of CHO cells. Metabolic Engineering. 23, 9–21 (2014).

40. A. Gouzy, G. Larrouy-Maumus, T. di Wu, A. Peixoto, F. Levillain, G. Lugo-Villarino, J. L. Gerquin-Kern, L. P. S. de Carvalho, Y. Poquet, O. Neyrolles, Mycobacterium tuberculosis nitrogen assimilation and host colonization require aspartate. Nature Chemical Biology. 9, 674–676 (2013).

41. D. Bottai, F. Levillain, A. Dumas, A. Gouzy, C. de Chastellier, T. Wu, R. Poincloux, R. Brosch, D. Schnappinger, L. P. So, Y. Poquet, O. Neyrolles, Mycobacterium tuberculosis Exploits Asparagine to Assimilate Nitrogen and Resist Acid Stress during Infection. 10 (2014), doi:10.1371/journal.ppat.1003928.

42. K. Y. Rhee, L. P. S. de Carvalho, R. Bryk, S. Ehrt, J. Marrero, S. W. Park, D. Schnappinger, A. Venugopal, C. Nathan, Central carbon metabolism in Mycobacterium tuberculosis: An unexpected frontier. Trends in Microbiology. 19 (2011), pp. 307–314.

43. A. Serafini, L. Tan, S. Horswell, S. Howell, D. J. Greenwood, D. M. Hunt, M. D. Phan, M. Schembri, M. Monteleone, C. R. Montague, W. Britton, A. Garza-Garcia, A. P. Snijders, B. VanderVen, M. G. Gutierrez, N. P. West, L. P. S. de Carvalho, Mycobacterium tuberculosis requires glyoxylate shunt and reverse methylcitrate cycle for lactate and pyruvate metabolism. Molecular Microbiology. 112, 1284–1307 (2019).

44. A. K. Pandey, C. M. Sassetti, “Mycobacterial persistence requires the utilization of host cholesterol” (2008), (available at www.pnas.orgcgidoi10.1073pnas.0711159105).

45. A. Rizvi, A. Shankar, A. Chatterjee, T. H. More, T. Bose, D. F. Gomez-casati, Rewiring of Metabolic Network in Mycobacterium tuberculosis During Adaptation to Different Stresses. 10, 1–16 (2019).

46. J. Bi, Y. Wang, H. Yu, X. Qian, H. Wang, J. Liu, X. Zhang, Modulation of Central Carbon Metabolism by Acetylation of Isocitrate Lyase in Mycobacterium tuberculosis. Scientific Reports. 7 (2017), doi:10.1038/srep44826.

47. H. Eoh, K. Y. Rhee, Methylcitrate cycle defines the bactericidal essentiality of isocitrate lyase for survival of mycobacterium tuberculosis on fatty acids. Proceedings of the National Academy of Sciences of the United States of America. 111, 4976–4981 (2014).

48. M. Kohlstedt, C. Wittmann, GC-MS-based 13 C metabolic flux analysis resolves the parallel and cyclic glucose metabolism of Pseudomonas putida KT2440 and Pseudomonas aeruginosa PAO1. Metabolic Engineering. 54, 35–53 (2019).

49. P. Basu, N. Sandhu, A. Bhatt, A. Singh, R. Balhana, I. Gobe, N. A. Crowhurst, T. A. Mendum, L. Gao, J. L. Ward, M. H. Beale, J. McFadden, D. J. V. Beste, The anaplerotic node is essential for the intracellular survival of Mycobacterium tuberculosis. Journal of Biological Chemistry. 293, 5695–5704 (2018).

50. J. L. Gallant, A. J. Viljoen, P. D. van Helden, I. J. F. Wiid, Glutamate dehydrogenase is required by mycobacterium bovis BCG for resistance to cellular stress. PLoS ONE. 11 (2016), doi:10.1371/journal.pone.0147706.

51. P. Droste, K. Nöh, W. Wiechert, Omix - A visualization tool for metabolic networks with highest usability and customizability in focus. Chemie-Ingenieur-Technik. 85, 849–862 (2013).

52. K. Nöh, P. Droste, W. Wiechert, Visual workflows for 13C-metabolic flux analysis. Bioinformatics. 31, 346–354 (2015).

53. D. J. V. Beste, T. Hooper, G. Stewart, B. Bonde, C. Avignone-Rossa, M. E. Bushell, P. Wheeler, S. Klamt, A. M. Kierzek, J. McFadden, GSMN-TB: A web-based genome-scale network model of Mycobacterium tuberculosis metabolism. Genome Biology. 8 (2007), doi:10.1186/gb-2007-8-5-r89.

54. D. J. v Beste, J. Peters, T. Hooper, C. Avignone, M. E. Bushell, J. Mcfadden, Compiling a Molecular Inventory for. Society. 187, 1677–1684 (2005).

55. S. Heller, A. McNaught, S. Stein, D. Tchekhovskoi, I. Pletnev, InChI - The worldwide chemical structure identifier standard. Journal of Cheminformatics. 5 (2013), doi:10.1186/1758-2946-5-7.

56. M. Beyß, S. Azzouzi, M. Weitzel, W. Wiechert, K. Nöh, The design of FluxML: A universal modeling language for 13C metabolic flux analysis. Frontiers in Microbiology. 10 (2019), doi:10.3389/fmicb.2019.01022.

57. P. Millard, F. Letisse, S. Sokol, J. C. Portais, IsoCor: Correcting MS data in isotope labeling experiments. Bioinformatics. 28, 1294–1296 (2012).

58. M. Weitzel, K. Nöh, T. Dalman, S. Niedenführ, B. Stute, W. Wiechert, 13CFLUX2 - High-performance software suite for 13C-metabolic flux analysis. Bioinformatics. 29, 143–145 (2013).

59. M. Weitzel, W. Wiechert, K. Nöh, The topology of metabolic isotope labeling networks. BMC Bioinformatics. 8 (2007), doi:10.1186/1471-2105-8-315.

60. J. F. Jadebeck, A. Theorell, S. Leweke, K. Nöh, HOPS: high-performance library for (non-)uniform sampling of convex-constrained models. Bioinformatics (Oxford, England). 37, 1776–1777 (2021).

61. A. Theorell, J. F. Jadebeck, K. Nöh, J. Stelling, PolyRound: polytope rounding for random sampling in metabolic networks, Bioinformatics. 38, 566–567 (2022).

62. A. Gelman, D.B. Rubin. Inference from iterative simulation using multiple sequences. Stat.Sci. 7, 457–472. (1992).

63. V. A. López-Agudelo, T. A. Mendum, E. Laing, H. H. Wu, A. Baena, L. F. Barrera, D. J. V. Beste, R. Rios-Estepa, A systematic evaluation of mycobacterium tuberculosis genome-scale metabolic networks. PLoS Computational Biology. 16, e1007533 (2020).

64. J. Beau, W. Webber, “A Bi-Symmetric Log transformation for wide-range data.”

65. W. R. Gilks, S. Richardson, D. Spiegelhalter, Eds., Markov Chain Monte Carlo in Practice: Interdisciplinary Statistics (Chapman & Hall/CRC Interdisciplinary Statistics Series), Chapman & Hall/CRC, 1st ed., 1995.

